# Brain histone beta-hydroxy-butyrylation couples metabolism with gene expression

**DOI:** 10.1101/2021.06.28.449924

**Authors:** Sara Cornuti, Siwei Chen, Leonardo Lupori, Francesco Finamore, Muntaha Samad, Francesco Raimondi, Raffaele Mazziotti, Christophe Magnan, Silvia Rocchiccioli, Pierre Baldi, Paola Tognini

## Abstract

Little is known about the impact of metabolic stimuli on brain tissue at a molecular level. The ketone body beta-hydroxybutyrate (BHB) can be a signaling molecule regulating gene transcription. Thus, we assessed lysine beta-hydroxybutyrylation (K-bhb) levels in proteins extracted from the cerebral cortex of mice undergoing a ketogenic metabolic challenge (48 hrs fasting). We found that fasting enhanced K-bhb in a variety of proteins including histone H3. ChIP-seq experiments showed that K9 beta-hydroxybutyrylation of H3 (H3K9-bhb) was significantly enriched by fasting on more than 8000 DNA loci. Transcriptomic analysis showed that H3K9-bhb on enhancers and promoters correlated with active gene expression. One of the most enriched functional annotations both at the epigenetic and transcriptional level was “circadian rhythms’’. Indeed, we found that the diurnal oscillation of specific transcripts was modulated by fasting at distinct zeitgebers both in the cortex and suprachiasmatic nucleus. Moreover, specific changes in locomotor activity daily features were observed during re-feeding after 48-hour fasting.

Thus, our results suggest that fasting dramatically impinges on the cerebral cortex transcriptional and epigenetic landscape, and BHB acts as a powerful epigenetic molecule in the brain through direct and specific histone marks remodeling in neural tissue cells.

## INTRODUCTION

Nutrition is a key regulator of whole-body physiology and health. Food impinges on tissue functions through modulation of metabolism, which results in adaptive changes in cellular biochemical/molecular processes, and ultimately tissue homeostasis. Notably, besides the well-documented influence on peripheral organs’ physiology/pathology, mounting evidence shows that the metabolic status can also affect neuronal function and metabolism, and influence brain physiology and ultimately cognitive processes, emotions and behavior (Gómez-Pinilla, 2008; Mattson et al., 2018; Padamsey et al., 2022; Tognini et al., 2020). Indeed, the high fat/high sugar western diet negatively affects brain health, leading to alterations in cognitive functions, anxiety, depression, and in general a higher incidence of emotional disorders (Dixon et al., 2013; Dutheil et al., 2016). On the other hand, caloric restriction has been shown to prevent age-related brain damage and to be neuroprotective by influencing free radical metabolism, and the cellular stress response system (Luo et al., 2017; Walsh et al., 2014). In 1920’s very low carbohydrate/high fat ketogenic diets (KD) were introduced in the clinic to treat refractory epilepsy, mimicking the increase in the blood level of ketone bodies (i.e. beta-hydroxybutyrate (BHB)), which were thought to mediate the effect of fasting on seizure events (Lutas and Yellen, 2013). Notably, epilepsy has high comorbidity with neurodevelopmental disorders such as autism spectrum disorders (ASD), and KD is one of the therapeutic approaches proposed to treat ASD patients (Evangeliou et al., 2003; Li et al., 2021). KD seems also to be beneficial in the treatment of neurodegenerative and psychiatric diseases (Louw et al., 2016; Murakami and Tognini, 2022; Paoli et al., 2013). Finally, fasting and aerobic exercise may improve cognitive performance and protect against depression and anxiety-like behavior through the enhancement of hippocampal neurogenesis (Landry and Huang, 2021; Lee et al., 2002; Vivar et al., 2012). Thus, in short, diet robustly influences neurological outcomes, yet the molecular and biochemical underpinnings of these effects are poorly understood (Pizzorusso and Tognini, 2020).

KD, fasting or starvation, and prolonged aerobic exercise, all promote the depletion of liver glycogen stores, favoring the decrease of blood glucose concentration. Due to its scarcity, glucose cannot be used to respond to the energy request of organismal tissues. Thus, to maintain metabolic homeostasis, our body switches to an alternative fuel: ketone bodies, produced via the activation of the so-called ketogenic pathway. Mainly in the liver, the acetyl-CoA produced by fatty acid oxidation is converted into three ketone bodies: acetoacetate, acetone, and BHB. The latter is the most abundant circulating ketone body and, under ketogenic conditions, the major energy source for metabolic active tissues, such as the brain (Newman and Verdin, 2014). Intriguingly, BHB is not just an energy carrier as it was believed in the past. Emerging evidence suggests that BHB is also a signaling molecule, capable of binding to the HCAR2 and FFAR3 receptors (Kimura et al., 2011; Taggart et al., 2005; Won et al., 2013), and of acting as an endogenous epigenetic regulator. Indeed, BHB selectively acts as a class 1 Histone deacetylase inhibitor, increasing histone acetylation and active gene expression (Shimazu et al., 2013; Tognini et al., 2017). Finally, BHB is the chemical donor for a new epigenetic mark so far investigated in detail only in the liver: lysine beta-hydroxy-butyrylation (K-bhb). Liver K-bhb contributes to active gene expression and regulates genes involved in the hepatic metabolic response to starvation (Xie et al., 2016).

Despite the high ketone bodies demand of the brain, the impact of BHB on neural tissue epigenome and transcriptome has never been explored. To address this important issue, which could underline and shed light on the adaptation capability of neuronal circuits to nutritional changes, we analyzed *in vivo* neural tissue responses to a specific metabolic challenge consisting of 48-hour fasting. Specifically, we discovered that fasting enhanced K-bhb in a variety of proteins. Then, we explored the genome-wide distribution of K9-bhb on histone H3 (H3K9-bhb) through chromatin immunoprecipitation, followed by sequencing (ChIP-seq) in the occipital cortex (an area corresponding to the visual cortex) of fasted (F) mice, and parallel measurement of the transcriptional response through RNA-seq. Dataset crossing pointed toward H3K9-bhb being an epigenetic mark related to fasting-induced gene expression, therefore suggesting that fasting-driven increase in BHB could directly affect brain cells through the remodeling of the chromatin landscape and transcriptional responses.

## RESULTS

### Fasting driven-increase in BHB causes a burst in protein Lysine beta-hydroxybutyrylation in the brain

To assess the molecular response of brain tissue to fasting-induced ketosis, we adopted a protocol of prolonged fasting and we assessed the biochemical action of a major metabolite produced upon fasting: BHB (Fig. 1A). Postnatal day (P) 33 mice were subjected to fasting (F) condition, corresponding to a total absence of food for 48 hours (48h), but ad libitum (AL) access to water. Their body weight and glycemia were monitored before and after fasting. F mice displayed a significant decrease in their body weight and blood glucose concentration after 48h, while AL control mice did not show any significant difference (Fig. 1 B-C). During the 48 h of food deprivation, F animals displayed a significant increase in spontaneous locomotor activity both during night and day time (Fig. 1D), showing that our protocol did not cause lethargy in the subjects and confirming behavioral changes reflecting the search for food (Koubi et al., 1991). Since BHB has been demonstrated to be the chemical moiety for a new post-translational modification (PTM) on K residues (Koronowski et al., 2021; Xie et al., 2016), we performed a western-blot analysis of protein extract from the occipital cortex (corresponding to the mouse visual cortex) of F and AL mice using a pan-K-bhb antibody. Strikingly, K-bhb was significantly increased in the occipital cortex of F mice with respect to AL (Fig. 2A), suggesting that the increased cortical BHB after fasting could be exploited, not only to produce energy, but also as a chemical donor for PTM. As a control, we analysed the pan-K-bhb in the liver of the same mice and confirmed a significant upregulation in F mice with respect to AL (Fig. S1A).

**Figure 1.**
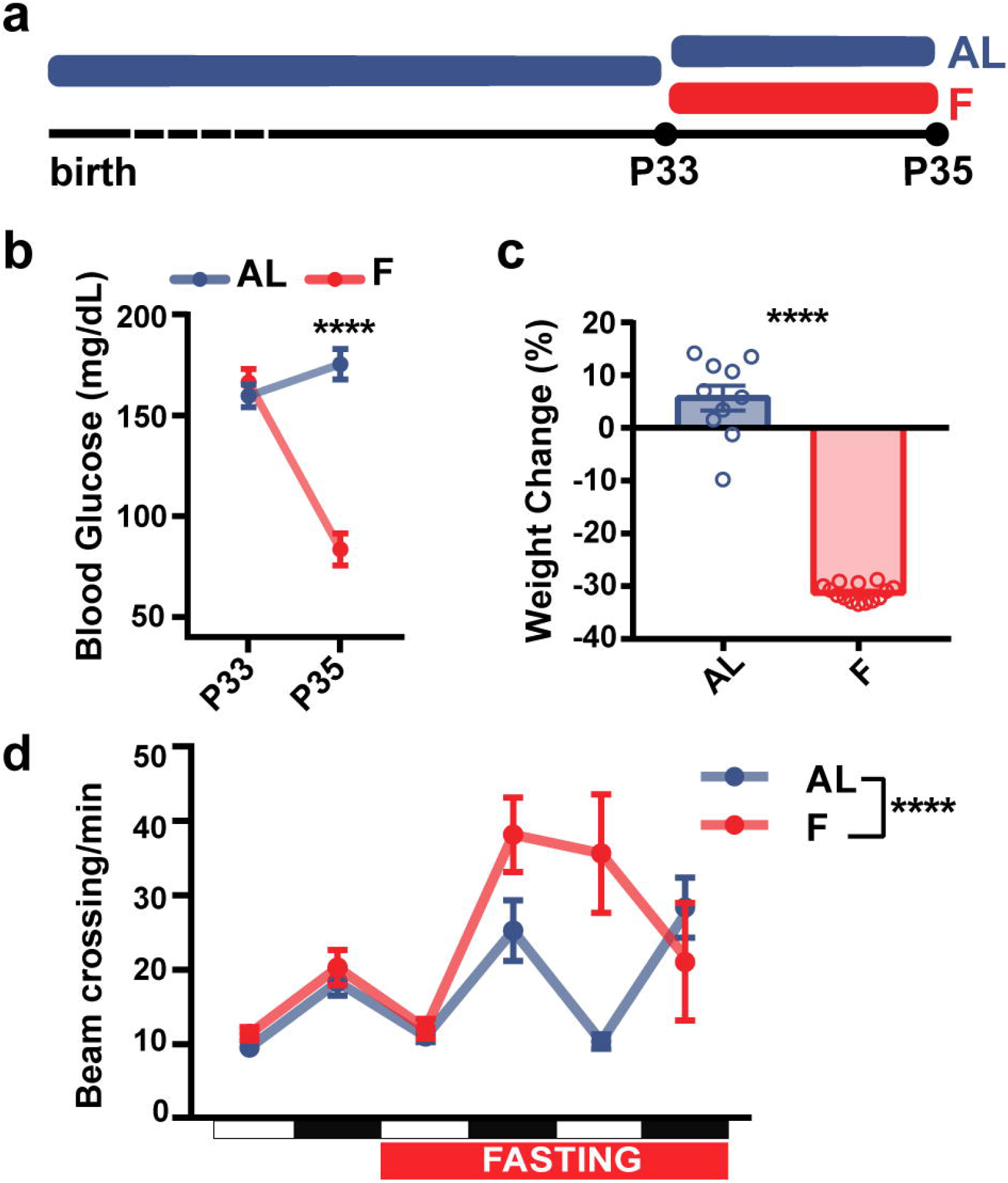
Fasting affects blood glucose concentration and locomotor activity. (**a**) Experimental timeline. (**b**) Blood glucose concentration (mg/dL) before and after 48h fasting (AL N=9, F N=10, two-way RM ANOVA time*treatment, interaction F_1,17_=64.19 p<0.0001, post-hoc Sidak, AL vs F (postFASTING) t_34_=9.203 p<0.0001); (**c**) Weight change (%) before and after 48h fasting (N=10 AL, N=14 F, unpaired 2-tailed t-test, t_22_=18.06 p<0.0001); (**d**) Spontaneous locomotor activity throughout the daily cycle before and during fasting (N=10 AL, N=7 F, two-way RM ANOVA time*treatment, interaction F_5,75_=6.596 p<0.0001).White and black squares on the x axes represent day-time and night-time respectively Error bars represent SEM.

HPLC-MS analysis of K-bhb residues in protein extract from the occipital cortex highlighted the presence of 234 beta-hydroxybutyrilated proteins. In total, 137 proteins were beta-hydroxybutyrilated exclusively in F mice, 77 proteins exclusively in AL mice, and 20 proteins were beta-hydroxybutyrilated in both groups (Fig. 2B). The 20 common beta-hydroxybutyrilated proteins were enriched in annotations such as “NADP metabolic processes” and “positive regulation of fibroblast proliferation”‘(Fig. 2B and Dataset S1). On the other hand, the F proteins displaying K-bhb clustered in specific GO annotations mainly related to “regulation of transcription, regulation of transcription-dependent on RNA pol II and morphogenesis” (Fig. 2B and Dataset S2). Also, the GO biological processes analysis of the proteins beta-hydroxybutyrilated in AL indicated an enrichment in “transcription, and chromatin modification” annotations (Fig. 2B, and Dataset S3). These findings suggest that BHB may have a more complex function related to transcriptional regulation, not just limited to HDAC inhibition or being a chemical moiety for histone PTMs, but also associated with the modulation of epigenetic (e.g. KDM1B, HDAC2), transcription factors (e.g. E2F4, HOXB3) or other chromatin-related proteins (e.g. SKOR1, CHD5) functions.

**Figure 2.**
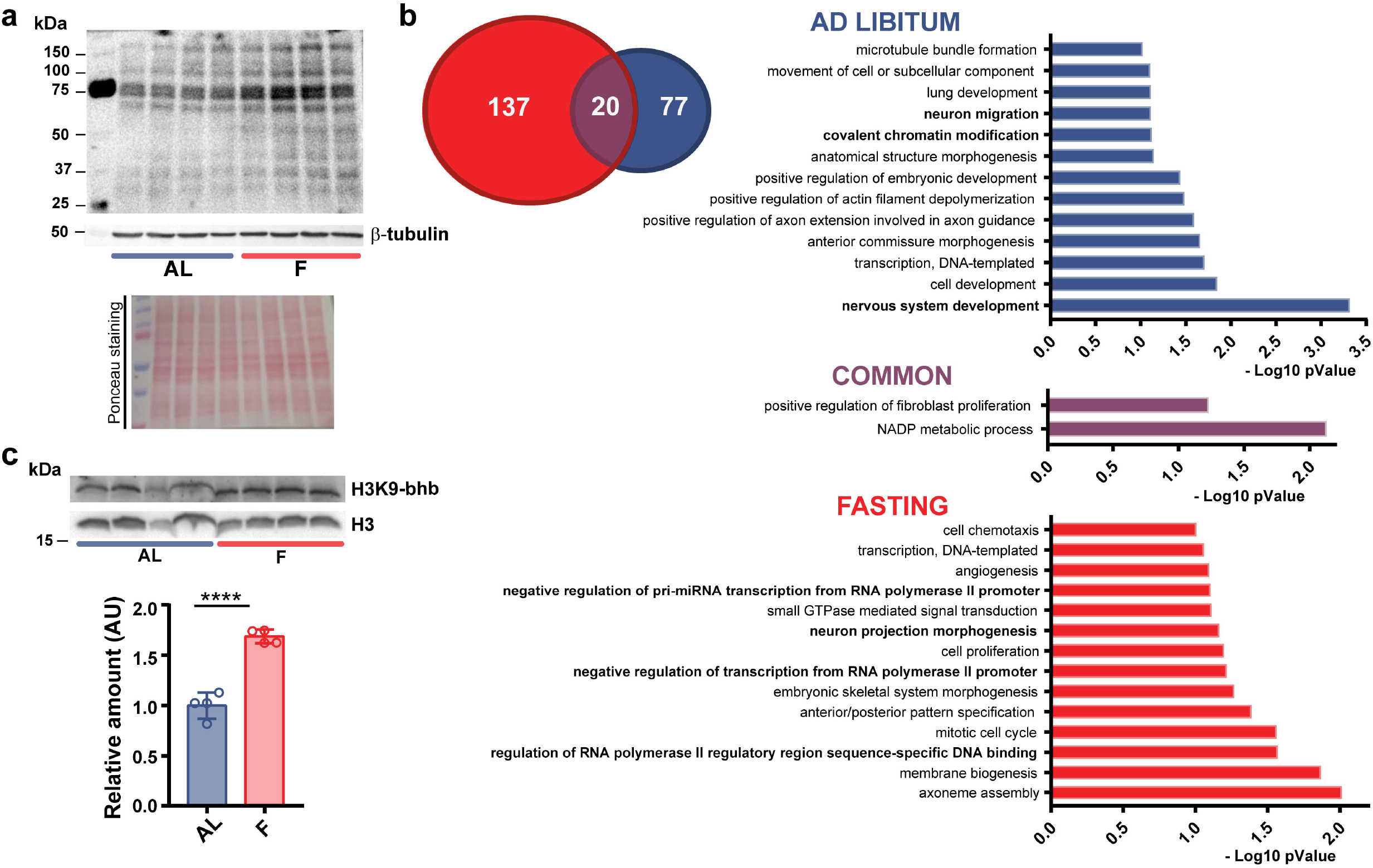
Fasting increases protein Lysine beta-hydroxybutyrylation and modifies the beta-hydroxybutyrylation profile of the total proteome in the cerebral cortex. (**a**) Western blot analysis of total lysine beta-hydroxybutyrylation in the occipital cortex of F or AL mice using a Pan-K-bhb antibody (top panel), a control beta-tubulin antibody (middle panel) and the Ponceau staining (bottom panel). (**b**) Left: Venn diagram of K-bhb proteins from the occipital cortex of AL (blue) and F mice (red). In purple the K-bhb proteins found in both conditions. Right: GO analysis (biological processes) of K-bhb proteins found in AL, F and in both groups (AL N= 3, F N= 3). (**c**) Western blot analysis of histone lysine beta-hydroxybutirylation in the occipital cortex of F or AL using a specific H3K9-bhb antibody (top panel) or a control H3 antibody (middle panel). Histograms representing the normalized ratio of H3K9-bhb /H3 in F condition compared to AL (AL N=4, F N=4, unpaired 2-tailed t-test, t_6_=9.361 p<0.0001). Error bars represent SEM.

BHB has been demonstrated to modify K-bhb on histones with consequent alteration of the liver transcriptional program (Xie et al., 2016). Thus, we studied K9-bhb on histone H3 (H3K9-bhb) in the brain using a specific antibody. H3K9-bhb was significantly more abundant in the cortex of F mice (Fig. 2C) with respect to the AL group. As expected, H3K9-bhb increased also in the liver of F mice (Fig. S1B).

Overall, these data demonstrate that beta-hydroxybutyrylation is conspicuously enhanced by fasting in neural tissue.

### BHB impacts the chromatin state of the cerebral cortex through a direct epigenetic action

To further explore the role of H3K9-bhb in the CNS, we performed a ChIP-seq experiment and studied the genome-wide distribution of H3K9-bhb in the occipital cortex of mice subjected to 48h fasting or fed AL. The ChIP-seq analysis revealed a striking increase in H3K9-bhb enriched loci in F with respect to AL. Indeed, the differential analysis between the two experimental conditions detected about 8400 enriched peaks (p<0.05) in F vs AL (Dataset S4) distributed in intergenic (42%), promoter (17%), and enhancer regions (18%, Fig. 3A). Importantly, only one peak was significantly enriched in AL with respect to F (Dataset S4), confirming that the increase in BHB observed during fasting is responsible for the dramatic increase in neural tissue H3K9-bhb.

**Figure 3.**
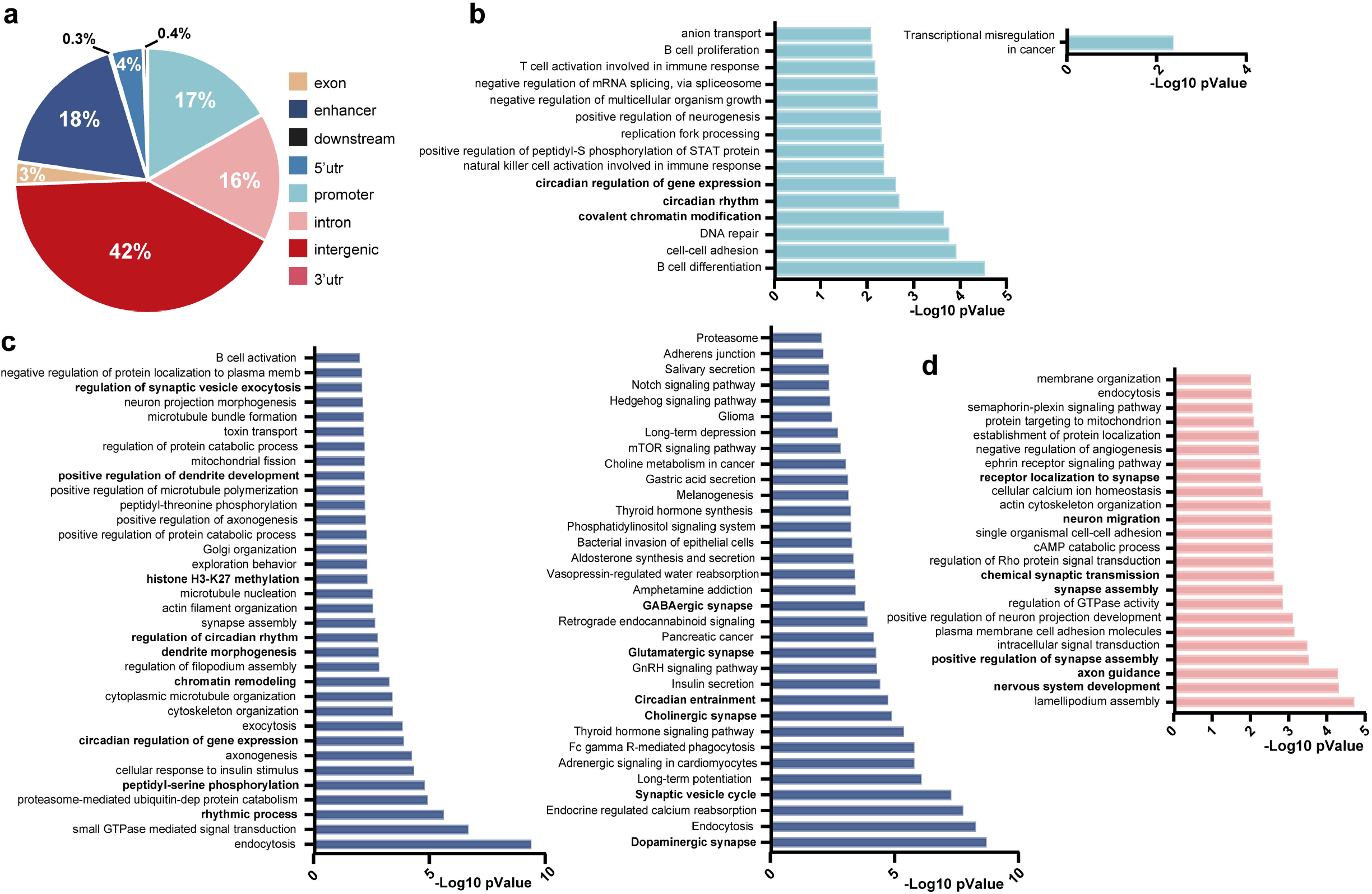
Fasting induces robust changes in H3K9 beta-hydroxybutyrylation in the mouse cerebral cortex. (**a**) Pie chart of genome-wide distribution of H3K9-bhb enriched loci in F with respect to AL (Cyber-Tp<0.05). (**b**) GO Biological processes (left graph) and KEGG pathway (right graph) analysis of the genes in proximity of promoters. (**c**) Same as b for enhancer regions. (**d**) Same as b for introns. No significant KEGG pathway was found for gene introns.

To gain further insight into the specific epigenetic effects of BHB in neural tissue, we crossed our H3K9-bhb ChIP-seq data with the already published H3K9-bhb ChIP-seq data in the liver (Xie et al., 2016). The analysis of the data in Xie et al., with our pipeline (see materials and methods) revealed that the total number of H3K9-bhb peaks in the brain was lower than in the liver (8404 brain loci vs 15416 liver loci). Peak category distribution demonstrated that H3K9-bhb was principally present in promoter and enhancer regions in the mouse liver after 48h fasting (Fig. S2), while enhancer, promoter, and especially intergenic regions were the most prominent category in the brain (Fig. 3A). In the cerebral cortex, the GO “biological processes” analysis of the genes in proximity of promoters and enhancer regions (Datasets S5 and S6) highlighted a specific signature related to circadian rhythms, and pathways involving histone PTM and chromatin remodeling (Fig. 3B-C). Also, synaptic transmission, dendrite morphogenesis, and synapse assembly came up as enriched GO terms associated with the enhancer regions. Brain neurogenesis, brain and dendrite morphogenesis, and axon guidance terms were dominant among the genes associated with the promoter regions (Dataset S5). The KEGG pathway analysis in the enhancers confirmed the “circadian rhythms” and “synaptic transmission” categories (Fig. 3C, Dataset S6). Finally, intergenic regions showed enriched annotations about synapse regulation, transmission and plasticity (Fig. 3D, Dataset S7).

Data crossing revealed that H3K9-bhb was present in 547 promoter regions and 681 enhancers common to brain and liver (Dataset S8). KEGG pathway analysis highlighted several enriched annotations related to metabolism for the common enhancers, among them “Insulin signaling”, “mTOR signaling”, “FoxO signaling” pathway, and “circadian entrainment” and “circadian rhythms” (Dataset S9). “FoxO signaling” pathway was present also in the common promoter list (Dataset S10).

Overall, these data demonstrate that 48h fasting is able to remodel the neural chromatin landscape, inducing a dynamic and robust H3K9-bhb in the cerebral cortex, preferentially targeting enhancers, promoters and intergenic regions.

### H3K9-bhb is linked to active gene expression in neural tissue

To investigate the impact of H3K9-bhb on gene expression in the visual cortex, we performed an RNA-seq experiment using the same cortical samples used for the ChIP-seq experiment. Fasting dramatically remodeled the transcriptome of the cortex altering the expression of 955 transcripts (Benjamini-Hochberg <0.01. For the complete gene list see Dataset S11).

In particular, 341 genes were upregulated and 614 genes were downregulated in F mice (Fig. 4A). The effect was highly consistent in the different samples, as shown by the heatmap obtained from every biological replicate (Fig. 4A). The most overrepresented KEGG pathways among the genes downregulated in fasting were “steroid biosynthesis”, “protein processing in endoplasmic reticulum”, “metabolic pathway”, “purine metabolism” etc. (Fig. 4B, Dataset S12A); categories mainly related to metabolism, thus suggesting the existence of a brain adaptive response to conserve energy. KEGG pathways characterizing the UP in fasting transcripts were “serotonergic synapses”, “oxytocin signaling pathway”, and “p53 signaling pathway” (Fig. 4B, Dataset S12B). Notably, the most significant KEGG pathway in the genes upregulated in the neocortex after fasting was “circadian rhythms’’ (Fig. 4B); this is of interest since core-clock genes have been involved in the control of critical period onset in the mouse visual cortex (Kobayashi et al., 2015). Finally, the interaction network of the 955 differentially expressed genes revealed a variety of interactomes including “FoxO” “mTOR” “Insulin” and “PPAR” signaling pathways which are well-known biochemical cascades influenced by fasting; “Glycosaminoglycan metabolism” “Chondroitin sulfate metabolism and biosynthesis” which are components of the perineuronal nets, important for ocular dominance plasticity and critical period timing (Pizzorusso et al., 2002); and again “circadian rhythms” (Fig. S3; for a complete list see Dataset S13).

**Figure 4.**
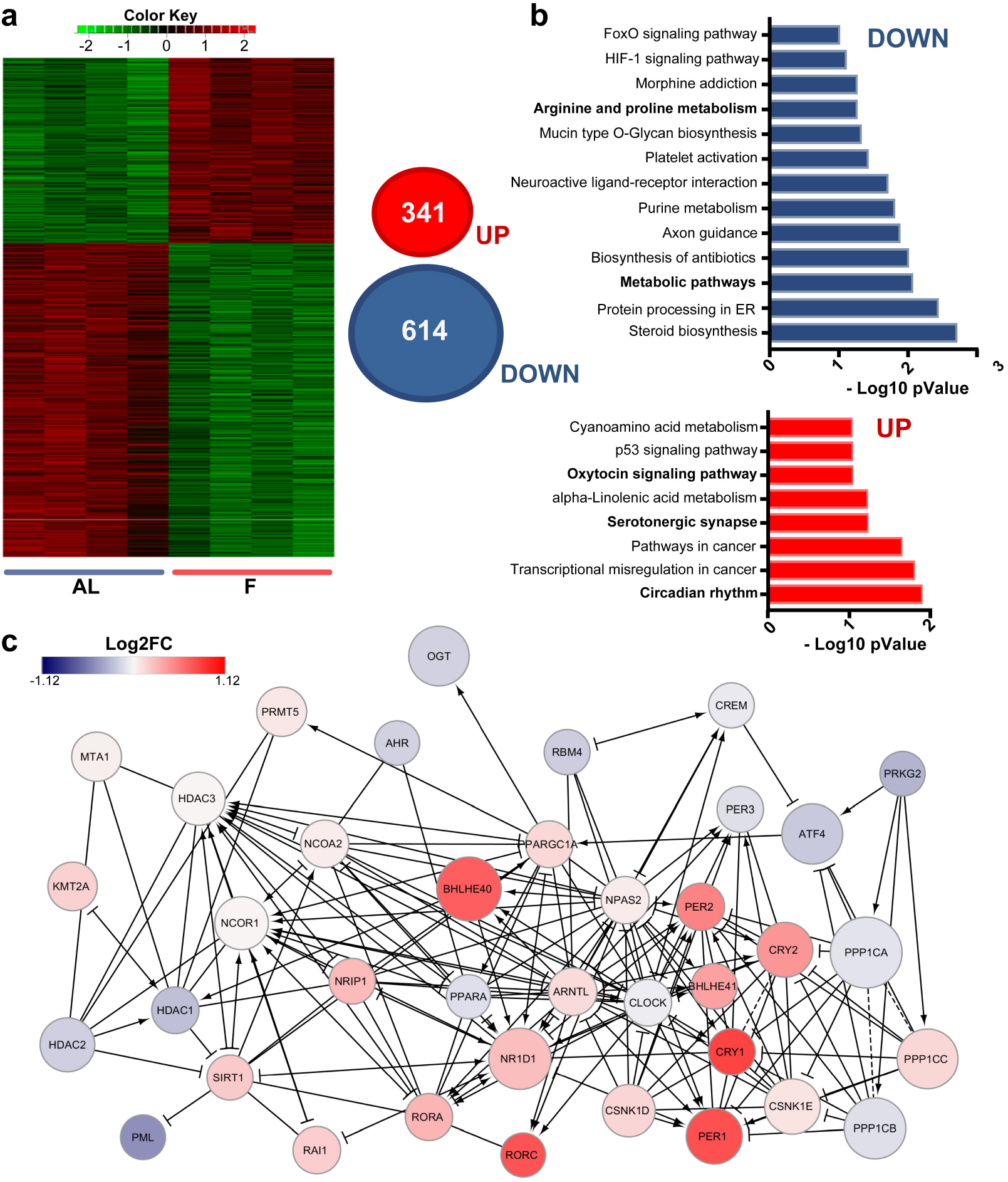
Fasting is related to active gene expression in the cortex of young mice. (**a**) Left: Heat map of RNA-seq analysis performed in the occipital cortex of F and AL mice (AL N=4, F N=4, p<0.05). Right: Venn diagrams representing the differential expressed genes (DEGs) between AL and F in the RNA-seq dataset (n=4, Benjamini-hochberg<0.01). (**b**) KEGG pathways analysis of DEGs in the occipital cortex of F mice. Upper panel: KEGG analysis of DEGs downregulated in the F condition. Lower panel: KEGG analysis of DEGs upregulated in the F condition. (**c**) Reactome pathway of DEG enriched in “circadian rhythms” annotation. Colors: Log2FC. Circle dimensions: Fasting means.

Finally, gene set enrichment analysis on the whole RNA-seq data set using CAMERA (Wu and Smyth, 2012) confirmed that “circadian rhythms” is significantly enriched in both GO and KEGG ontologies (Fig. S4A-B). Intriguingly, the Reactome pathway displayed a relationship between “circadian rhythms” terms and a variety of proteins related to epigenetic remodeling (Fig. 4C), suggesting that an interplay between circadian and epigenetic mechanisms could be responsible for the brain adaptive response to prolonged fasting.

To understand the role played by H3K9-bhb on the cerebral cortex transcriptome, we crossed our RNA-seq and ChIP-seq data. We found a significant positive correlation between differential H3K9-bhb peaks (F vs AL) and the fold change of the genes upregulated in F (Fig. 5A, all the genes of the RNA-seq up in fasting, Spearman correlation: rho = 0.13, p value = 4.4E-16, only genes up in fasting with p value < 0.05, Spearman correlation: rho = 0.19, p value = 6.7E-09). Furthermore, the expression of genes in proximity of loci differentially enriched in H3K9-bhb peaks was selectively upregulated by fasting with respect to all expressed genes (Fig. 5B, Mann-Whitney test: U = 19670177, p < 0.0001). It is worth noting that there was no significant correlation between the differential H3K9-bhb peaks and the genes upregulated in AL condition (Fig. 5C, all the genes of the RNA-seq up in AL, Spearman correlation: rho = 0.0008, p value = 0.96. Fig. 5C only genes up in AL with p value < 0.05, Spearman correlation: rho = -0.02, p value = 0.41). These results suggest that H3K9-bhb could be related to fasting-induced gene expression in the cerebral cortex of mice.

**Figure 5.**
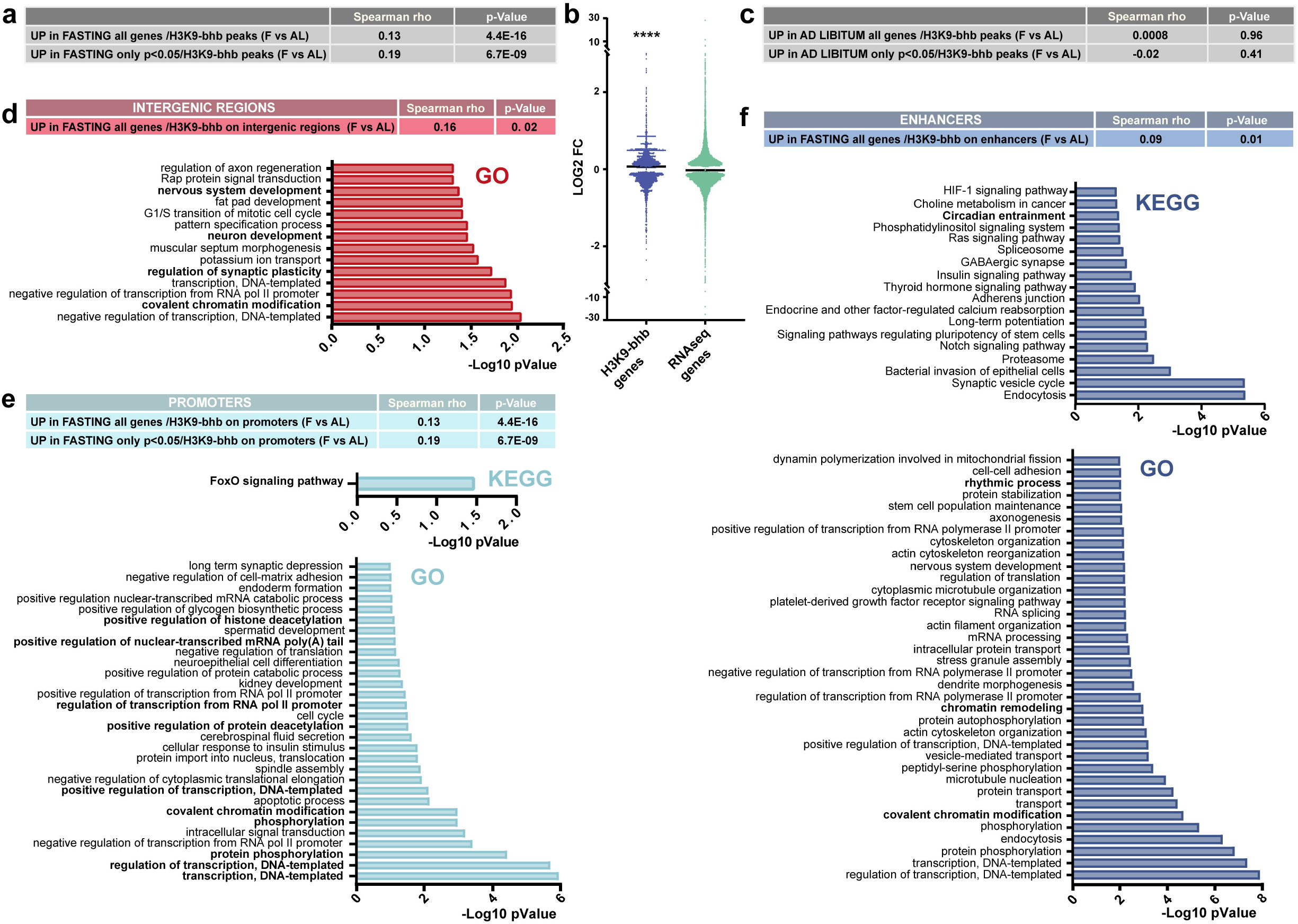
H3K9-bhb is linked to active gene expression in the cortex of fasted mice. (**a**) Correlation between F-induced H3K9-bhb peak enrichment and transcript fold change revealed by RNA-seq (F vs AL, all the genes of the RNA-seq up in fasting, Spearman correlation: rho = 0.13, p value = 4.4E-16; only genes up in fasting with p value < 0.05, Spearman correlation: rho = 0.19, p value = 6.7E-09). (**b**) Fold change expression induced by fasting is significantly higher for genes in proximity to H3K9-bhb peaks than for all genes. Horizontal bar represents median, Mann-Whitney test: U = 19670177, p < 0.0001. (**c**) Lack of correlation between AL-induced H3K9-bhb peak enrichment and transcript fold changes revealed by RNA-seq (all the genes of the RNA-seq up in AL, Spearman correlation: rho = 0.0008, p = 0.96; only genes up in AL with p < 0.05, Spearman correlation: rho = -0.02, p = 0.41). (**d**) Top: Correlation between H3K9-bhb enriched intergenic regions and transcripts upregulated in fasting (only genes up in fasting with p < 0.05, Spearman correlation: rho = 0.16, p = 0.02). (**e**) Top: Correlation between H3K9-bhb enrichment on promoters and transcript fold-change for genes upregulated in fasting (all the genes of the RNA-seq up in fasting, Spearman correlation: rho = 0.17, p = 5.4E-08; only genes up in fasting with p value < 0.05, Spearman correlation: rho = 0.32, p = 1.2E-07). Bottom: KEGG pathways analysis and GO analysis of the H3K9-bhb promoters of genes upregulated in the fasting condition. (**f**) Top: Correlation between H3K9-bhb enrichment on enhancers and transcript fold-change for genes upregulated in fasting (all the genes of the RNA-seq up in fasting, Spearman correlation: rho = 0.09, p = 0.01). Bottom: KEGG pathways analysis and GO analysis of the H3K9-bhb enhancers of genes upregulated in the fasting condition.

To gain a better insight into the transcriptional regulatory role of cortical K-bhb after 48h fasting, we analyzed the list of differential and significantly enriched H3K9-bhb DNA loci (Dataset S4) separately considering promoters, enhancers and intergenic regions, and crossed them with the correspondent transcript levels obtained through the RNA-seq. H3K9-bhb was particularly abundant in intergenic regions (Fig. 3A). Epigenetic marks present in intergenic regions of brain chromatin have been shown to be sensitive to changes in neuronal activity, such as DNA methylation (Guo et al., 2011). Fasting-driven changes in H3K9-bhb in intergenic regions positively correlated with the expression of correspondent transcripts upregulated in fasting (Fig. 5D, only genes up in fasting with p value < 0.05, Spearman correlation: rho = 0.16, p value = 0.02). GO biological processes analysis revealed the association with genes involved in regulation of transcription, chromatin modifications, synaptic plasticity and neurodevelopment (Fig. 5D). Promoters and enhancers are key elements for controlling transcripts’ levels, and they were especially influenced by H3K9-bhb in F mice (Fig. 3A). We found a significant correlation between promoter hits and transcripts upregulated in fasting (Fig. 5E, all the genes of the RNA-seq up in fasting, Spearman correlation: rho = 0.17, p value = 5.4E-08, only genes up in fasting with p value < 0.05, Spearman correlation: rho = 0.32, p value = 1.2E-07). The genes clustered in the “FoxO signaling pathway” (Fig. 5E). Moreover, GO analysis of the term “Biological Process”, showed a variety of annotations principally related to transcriptional regulation and PTMs (Dataset S14).

The correlation of RNA-seq genes with H3K9-bhb enhancers gave again a positive value (Fig. 5F, all the genes of the RNA-seq up in fasting, Spearman correlation: rho = 0.09, p value = 0.01). Interestingly, pathways and terms concerning “circadian rhythms” came up both in the KEGG (Fig. 5F and Dataset S15) analysis and GO annotation (Fig. 5F and Dataset S15). Furthermore, GO biological processes related to “chromatin remodeling and modifications” were significant terms. Again and in keeping with the above analysis, we observed the presence of a signature specifically related to circadian clock and epigenetic mechanism, strongly suggesting that those could be the molecular underpinning underlying the cerebral cortex adaptation to a fasting challenge.

### Fasting influences diurnal gene expression of core-clock genes and daily rhythms in locomotor activity

Protein synthesis regulation and circadian rhythms are important mechanisms in neural function and plasticity (Hartsock and Spencer, 2020; Martin et al., 2000). Since GO categories associated with these mechanisms consistently emerged in our RNA-seq and ChIP-seq data, we selected these two pathways for in depth analysis of fasting regulation. First, we studied the “insulin and mTOR pathways” by assessing two key biochemical steps: serine 235-36 phosphorylation of S6 (Suppl. Fig. 5A) and Serine 473 phosphorylation of AKT (Fig. S5B). Confirming the hypothesis indicated by the RNA- and ChIP-seq data, both PTMs were significantly reduced in fasting mice, suggesting that protein synthesis could be altered in the cerebral cortex of F mice, in line with a condition of energy saving. Second, we studied “circadian rhythms”, a process representing one of the top hits in the RNA-seq data (Fig. 4B) and in the differential ChIP-seq enhancer analysis (Fig. 5E). We analyzed the expression of core-clock genes in the visual cortex (Fig. 6A) of F and AL mice at 4 different time points (Zeitgeber time (ZT)) every 6 hours throughout the daily cycle (Fig. 6B). The results demonstrated that several clock genes related to transcriptional inhibition: *Per1, Cry1, Cry2*, and *Reverb-a*, displayed similar changes in their diurnal profile upon fasting, and were significantly upregulated at ZT4 in F with respect to AL mice, validating the RNA-seq result which was performed at this ZT (Fig. 6C). On the other hand, *Bmal1, Clock* and *Ror-a*, belonging to the transcriptional activation limb of the core-clock machinery, were not significantly altered at any ZT in F mice (Fig. 6D), suggesting a certain degree of specificity of fasting effects on the inhibitory limb of the molecular clock. Prompted by these findings obtained in the visual cortex, we investigated whether circadian clock gene expression was also changed by fasting in the key brain structure for circadian rhythms, i.e. the suprachiasmatic nucleus (SCN) (Fig. S6A-B). We found that, as in the cortex, transcripts belonging to the negative limb of the core-clock were mainly impacted, with a significant increase at ZT22 (Fig. S6C). Moreover, *Bmal1* oscillation was strongly dampened (Fig. S6D).

**Figure 6.**
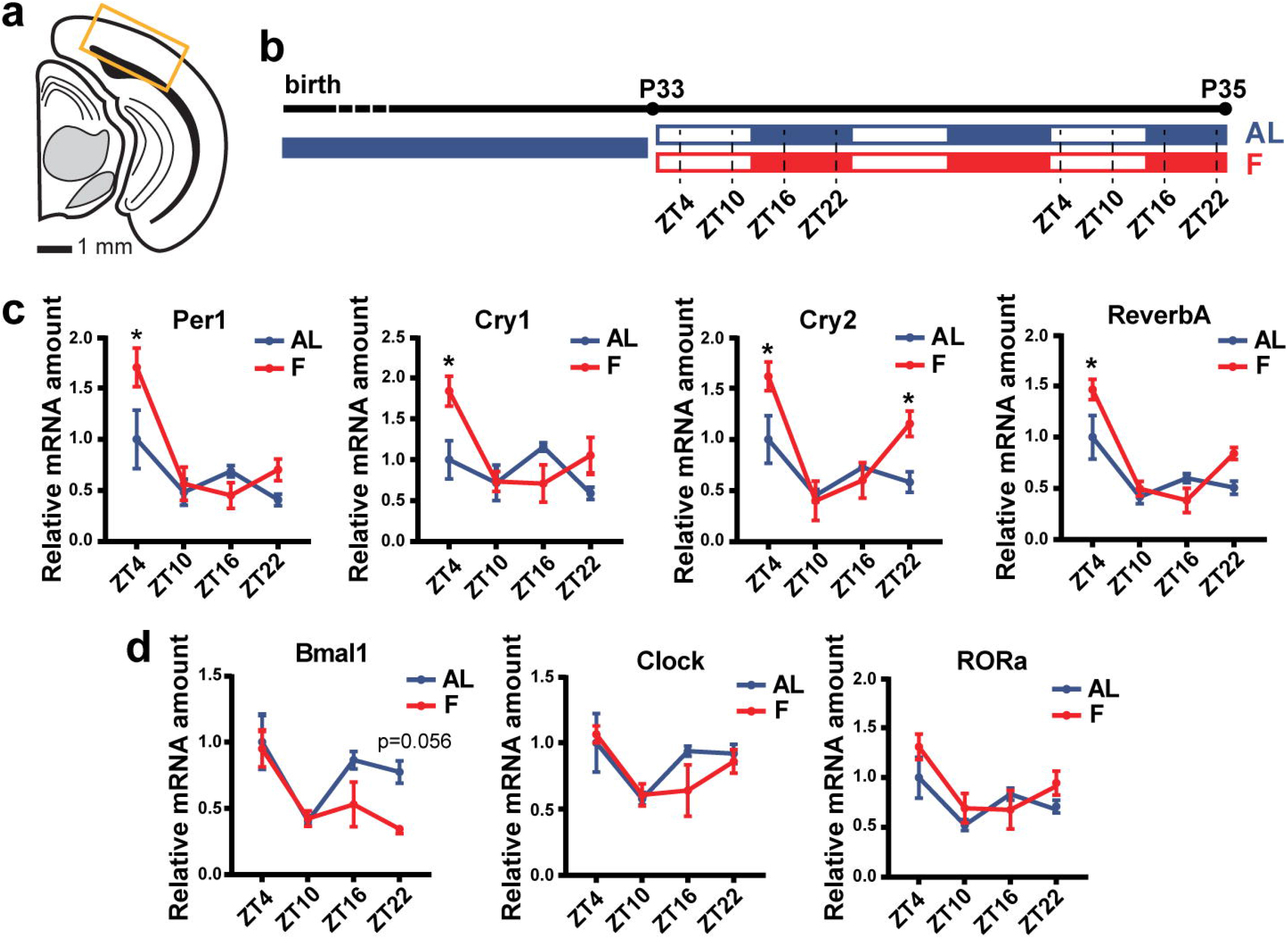
H3K9-bhb influences the expression of core-clock genes in the cortex of mice. (**a**) Schematic coronal section of the visual cortex showing the site of the gene expression analysis. (**b**) Experimental timeline. (**c**) Quantitative real-time PCR analysis of genes related to transcriptional inhibition of the core clock machinery (N=4 per condition, per time-point. Per1 :, two-way ANOVA time*treatment, interaction F_3,24_=3.234 p=0.04, post-hoc Sidak, AL vs F (ZT4) t_24_=3.209 p=0.015; Cry1: two-way ANOVA time*treatment, interaction F_3,24_=4.836 p=0.009, post-hoc Sidak, AL vs F (ZT4) t_24_=3.317 p=0.0115; Cry2: F, two-way ANOVA time*treatment, interaction F_3,24_=3.717 p=0.0251, post-hoc Sidak, AL vs F (ZT4) t_24_=2.999 p=0.0247, (ZT22) t_24_=2.769 p=0.042; Reverb-a, two-way ANOVA time*treatment, interaction F_3,24_=4.162 p=0.0165, post-hoc Sidak, AL vs F (ZT4) t_24_=3.144 p=0.0175). (**d**) Quantitative real-time PCR analysis of genes related to transcriptional activation limb of the core clock machinery (N=4 per condition, per time-point. Bmal1: F, two-way ANOVA time*treatment, interaction F_3,24_=1.167 p=0.1804; Clock: two-way RM ANOVA time*treatment, interaction F_3,24_=0.94 p=0.436; RORa: F, two-way RM ANOVA time*treatment, interaction F_3,24_=1.635 p=0.3869). Error bars represent SEM.

All these findings, together with the possible body-wide effect of fasting on circadian rhythmicity (Kinouchi et al., 2018; Vollmers et al., 2009), led us to explore whether fasting could alter the daily rhythmicity in spontaneous locomotor activity. Circadian rhythmicity in locomotion was investigated by selecting 3 different epochs of 5 days each: the 5 days before fasting (pre-fasting), the 5 days after refeeding (T1) and the subsequent 5 days (T2) (Fig. 7A). For every epoch, we computed the relative power spectrum at different oscillation frequencies and we integrated the power between 0.9 and 1.1 cycles/day as a measure of diurnal rhythmicity (Fig. 7B, see Material and Methods and Fig. S7B). The results demonstrated that before fasting there were no differences between AL and F groups (Fig. 7C). However, both at T1 and T2 (after fasting), F mice displayed a decreased power at the diurnal frequency (Fig. 7C), suggesting that acute fasting for 48h affects the normal development of rhythmicity in locomotion. Notably, this effect is persistent for at least 10 days after refeeding, and it could not be simply explained by alterations of neither total activity nor body weight (Fig. S7C, S7D), since both the measures showed no difference between groups both at T1 and at T2, thus suggesting that fasting might have a specific action on circadian rhythmicity.

**Figure 7.**
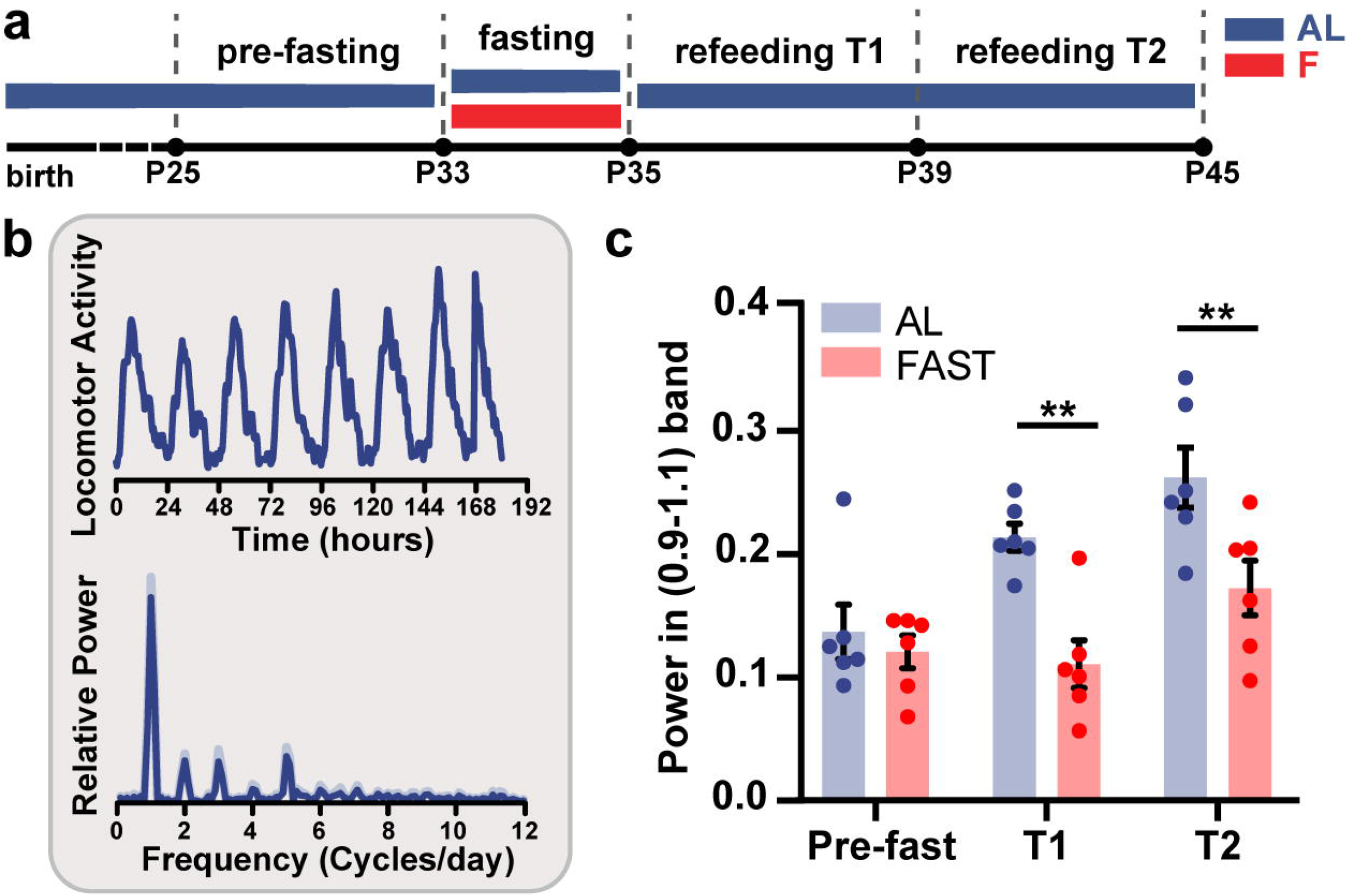
Fasting has long-lasting effects on the daily rhythmicity in spontaneous locomotor activity. (**a**) Experimental timeline. (**b**) Representative trace of the locomotor activity of a single mouse in the time domain (top) and in the frequency domain (bottom). (**c**) Relative power of the circadian oscillation (total power between 0.9 and 1.1 cycles per day) in AL and FAST mice in three epochs: before fasting (Pre-fast), first five days of refeeding (T1) and second 5 days of refeeding (T2). (N=6 per condition, two-way RM ANOVA time*treatment, interaction F_2,20_=4.11 p=0.032, post hoc Holm Sidak, AL vs FAST (Pre-fast) T_30_=0.598 p=0.555, AL vs FAST (T1) T_30_=3.768 p=0.002, AL vs FAST (T2) T_30_=3.274 p=0.005).

## DISCUSSION

Metabolic influence on the brain has been observed for years, however little is known about the molecular mechanisms activated by metabolic stimuli in brain tissue. In particular caloric restriction or intermittent fasting have been found to ameliorate brain resilience to aging processes (Mujica-Parodi et al., 2020), but its biochemical and transcriptional consequences on brain tissue in vivo are poorly understood. BHB, a metabolic fuel for the brain mainly produced by the liver during fasting, has been shown to be beneficial in models of neurological diseases (Yang et al., 2019), epilepsy (Simeone et al., 2017) and stroke (Stephan and Sleiman, 2021). These effects have been attributed to BDNF (Chen et al., 2017; Sleiman et al., 2016), HCAR2 (Rahman et al., 2014), and promotion of SIRT3 function (Yin et al., 2015). However, our study shows that a surprisingly high number of proteins can undergo K-bhb, raising the possibility that multiple unknown pathways can mediate BHB effects in the brain with potential therapeutic implications for brain disorders. To our knowledge, this is the first proteome-wide report of beta-hydroxybutyrylation in the cerebral cortex. A recent report investigated K-bhb in the liver after 48h fasting and found 267 beta-hydroxy butyrylated proteins, mainly included in several metabolic networks, energy and detoxification pathways (Koronowski et al., 2021). In the brain, the proteins displaying K-bhb belonged to specific GO categories which included regulation of transcription and chromatin modifications, raising the possibility of a cross-talk among beta-hydroxybutyrylation and other epigenetic mechanisms. Further studies are needed to investigate the BHB action on specific proteins and in the distinct cell types composing the neural tissue.

Our work also reveals a conspicuous transcriptomic change induced by fasting in cerebrocortical tissue, and demonstrates that BHB is an epigenetic molecule capable of dramatically remodeling the neural tissue chromatin landscape by post-translationally modifying the histone H3. Fasting-induced transcriptional changes and the enrichment of H3K9-bhb on a genome-wide scale were significantly correlated suggesting a regulatory role for H3K9-bhb in the neural tissue. Previous work studied H3K9-bhb response to the same fasting period used here on hepatic tissue (Xie et al., 2016). Our ChIP-seq analysis in cerebrocortical tissue revealed a number of DNA loci displaying differential H3K9-bhb after fasting summing up to about half of the DNA loci regulated in the liver, indicating that histone beta-hydroxybutyrylation may be a more pervasive PTM in liver than in neural tissue. Importantly, the data crossing of liver and cortex H3K9-bhb ChIP-seq data indicated several common promoters and enhancers among the two tissues. The KEGG pathways of the common enhancers were associated with “circadian rhythms” terms. The effects of fasting on the liver clock are well-known (Kinouchi et al., 2018; Vollmers et al., 2009), as well as the effects of H3K9-bhb on core-clock gene expression (Xie et al., 2016). However, how food deprivation could impinge on the cerebral cortex clock is still obscure. Our transcriptome and ChIP-seq data converge into a possible involvement of circadian processes in the brain response to a prolonged fasting (see below).

Based on genomic localization analysis, H3K9-bhb was mainly enriched in the cerebral cortex intergenic, enhancer and promoter regions of fasting mice. The genes neighboring those regions were positively correlated with the RNA-seq results of genes UP in fasting, extending and confirming previous findings in the liver (Xie et al., 2016), and suggesting that H3K9-bhb is a new PTM linked to active gene expression also in the brain. Although our understanding of intergenic regions function in the brain is still limited, H3K9-bhb in these genomic loci might help regulate the expression of associated transcribed regions, as already observed for DNA methylation in the human brain (Telese et al., 2013).

The functional annotation analysis of transcriptome, beta-hydroxybutyrylome and epigenome suggests that fasting could impinge on specific pathways, which could cooperate in setting up the best response of brain cells to a condition of scarce access to food. In particular, “circadian rhythms” is the most significantly enriched KEGG pathway among the upregulated genes, and “circadian entrainment”, “circadian regulation of gene expression”, “rhythmic process”, and again “circadian rhythms”, come up in the ChIP-seq functional annotation analysis of enhancer and promoter regions. Thus, diurnal rhythms seem to represent biological processes particularly impacted by fasting in the cerebral cortex. The circadian clock is an inner oscillator system that ensures appropriate physiology and fitness to a variety of organisms (Tognini et al., 2018; Zarrinpar et al., 2016). Changes in core-clock gene expression after 48h fasting might help to optimize the energetic cycle in neuronal cells, which consume much energy for electrical transmission and membrane potential maintenance; and/or non-neuronal cells, which need adequate fuel for surveillance/defense function (e.g.microglia), metabolism, throphism, and secretion/absorption of neurotransmitters, etc. (e.g. astrocytes). Intriguingly, our Reactome analysis of the cortical transcriptome displayed the existence of a link between circadian rhythms related genes and epigenetic related transcripts (Fig. 4C). Epigenetic mechanisms participate in the formation of circadian oscillation of gene expression, intricately regulating the core-clock machinery (Eckel-Mahan and Sassone-Corsi, 2013). The fasting challenge could directly control core-clock gene expression through a complex network of epigenetic proteins, possibly, fine tuning H3K9-bhb and other histone PTMs. Furthermore, the circadian clock controls the onset of critical period plasticity in the visual cortex (Kobayashi et al., 2015). Thus, we might speculate that the molecular/epigenetic/metabolic changes driven by fasting could contribute to altering the plasticity potential of the neural circuits through modulation of the endogenous 24h oscillator. An effect of fasting on plasticity is also in line with fasting modulation of protein synthesis, a regulatory mechanism involved in plasticity mechanisms (Cajigas et al., 2010), and the impact of the gut-brain axis on adult ocular dominance plasticity (Lupori et al., 2022). This hypothesis will be tested in future experiments.

Core-clock genes were also modulated in a time-of-the-day dependent manner in the SCN, suggesting the master clock could be sensitive to changes in the metabolic status (Tognini et al., 2020). As such, we found that fasting could interfere with normal rhythmicity in spontaneous locomotor activity (Fig. 7). The changes in rhythmicity of activity were observed both 5 days and 10 days after refeeding suggesting a long-lasting “memory trace” related to locomotion in F mice with respect to AL.

“Oxytocin signalling” and “serotonergic synapses” are categories present in our transcriptome functional annotation, and they seem particularly relevant for brain function. Oxytocin and serotonin are neuromodulators involved in human affects and socialization (Mottolese et al., 2014). For instance they have been shown to be implicated in ASD and depression (Borroto-Escuela et al., 2021; Yamasue and Domes, 2017). Prolonged fasting could promote social behaviour in order to cooperate and increase the probability to survive during periods of food scarcity (Weiss et al., 2017). Based on our GO results, intermittent fasting, caloric restriction, KD or ketone supplements might be alternative strategies or adjuvant to traditional treatments for ASD, depression or other neuropsychiatric diseases. This is an intriguing hypothesis which needs further validation in preclinical models to rationalize the use of nutrition to improve brain disorders, and cognitive function in general.

An important future step would be to deeply investigate the functional relevance of H3K9-bhb in the brain fasting response, and to causally link H3K9-bhb to fasting-induced gene expression. At this time, those experiments are particularly challenging since we have only little information about the mechanisms underlying the beta-hydroxybutyrylation chemical reaction. Recently, human SIRT3 was shown to display class-selective histone de-betahydroxybutyrylase activities on H3 K9-bhb (Zhang et al., 2019). SIRT3 seems to be preferentially a mitochondrial sirtuin (Verdin et al., 2010), while its nuclear localization is controversial (Bao et al., 2010; Cooper and Spelbrink, 2008; Hallows et al., 2008). Thus, genetically or pharmacologically modulating SIRT3 in the cortex to target H3K9-bhb during fasting could not be an appropriate approach based on SIRT3 uncertain subcellular localization and the unspecific effects caused by its deacetylase activity. Furthermore, Huang et al., discovered that the acyltransferase p300 can catalyze the enzymatic addition of BHB to K, while histone deacetylase 1 (HDAC1) and HDAC2 enzymatically remove K-bhb in cells in vitro (Huang et al., 2021). Since p300 is an acetyl-transferase, blocking its activity would be highly unspecific. A desirable approach could be to directly target H3K9-bhb in vivo for example through the use of an intrabody against this specific PTM (Chirichella et al., 2017), however this type of tool is not available yet.

In summary, our work shows that fasting is a powerful regulator of the transcriptional status of neural tissue modifying the chromatin epigenetic landscape leaving a trace on circadian behavior that can persist at the behavioral level. The regulated genes are not only involved in brain tissue metabolism, but they also include pathways relevant for synaptic transmission and plasticity, raising the hypothesis that the metabolic status of the individual might exert an important regulation on brain function.

## MATERIALS AND METHODS

### Animals and feeding

All experiments were carried out in accordance with the European Directives (2010/63/EU) and were approved by the Italian Ministry of Health (authorization number 354/2020-PR). C57BL/6J mice were used in this study. Mice were housed in conventional cages (365 × 207 × 140 mm, 2-3 animals per cage) with nesting material. Mice were kept under a 12 hour dark: 12 hour light cycle, with food (standard diet mucedola 4RF25) and water ad libitum.

Mice were weaned on postnatal day (P) 21. At P33 a group of mice was subjected to complete fasting (F group) for 2 days, until P35, through the removal of the food from the cage and ad libitum access to water. Age-matched control mice (AL group) continued to have ad libitum access to food and water. After two days, mice were sacrificed by decapitation. Liver and occipital cortices (corresponding to the visual cortex) were harvested, snapped frozen in liquid nitrogen and stored at -80 until use. For the day vs night sample analysis, mice were sacrificed every 6 hours along the diurnal cycle (from ZT0 to ZT24). For every group of mice, the food was precisely removed 48 hours before the ZT of the sacrifice.

### RNA sequencing and data analysis

N=4 biological replicates per experimental group were used in the RNA-seq experiment. Total RNA was extracted as described above, libraries were prepared and sequenced on Illumina HiSeq2500 instrument during a pair-end read 125bp sequencing, producing sequencing results in FastQ format. The FastQ files were processed through the standard Tuxedo protocol, using Tophat and Cufflinks. Tophat was used to align the RNA-seq reads to the reference genome assembly mm10 and Cufflinks was used to calculate gene expression levels. This protocol outputs the FPKM values for each gene of each replicate. The differential analysis of the FPKM values across all experiment and control groups was conducted with Cyber-T, a differential analysis program using a Bayesian-regularized t-test (Baldi and Long, 2001; Kayala and Baldi, 2012). The p-value threshold used for determining differential expression was 0.05 for all groups. The statistic was corrected using the Benjamini-Ochberg (BH) test for significance.

### ChIP-sequencing and analysis

N=4 biological replicates per experimental group were used for the sequencing. IPs and controls were processed in parallel and libraries were prepared at the Institute of Applied Genomics (Udine, Italy). The sequencing was performed on Hiseq2500 in 125 bp paired-end mode. The sequencing reads were aligned to the mouse genome assembly mm10 with Bowtie2. Enriched genome areas, and peaks, were identified with the MACS2 algorithm. Peak annotation was performed with in-house annotating tools. Peaks whose start to end position cover regions including promoter, enhancer, utr3, utr5, exon, intron, 2000 base pair downstream of promoter, or intergenic region were annotated and the percentages of the peak’s coverage of those regions were calculated. The closest gene and TSS to each peak were identified and their distances to the peak were calculated.

### Analysis of locomotor activity

Locomotor activity was measured using Opto M3 multi-channel activity monitors (Columbus Instruments, OH, USA). Monitors were placed in the colony area and testing was conducted in the same conditions of animal facility housing with the exception of animals being individually housed in (33 × 15 × 13) cm (length × width × height) clear plastic cages. Locomotor activity was measured by integrating the number of infrared beam breaks over 1-minute epochs for many days.

Total activity over an epoch was calculated as the sum of all beam brakes in that epoch. Diurnal rhythmicity was calculated by analyzing 5 days-long epochs in the frequency domain. For each mouse, we removed the non-oscillatory component of the activity trace by subtracting the mean activity, then we computed the periodogram power spectral density estimate at frequencies between 0.1 and 12 cycles/day in increments of 0.1. For each mouse, we normalized the power spectrum so that total power was equal to 1. The power of the circadian oscillation was defined as the sum of the normalized power between 0.9 and 1.1 cycles/day.

### Statistical analysis

The majority of statistical analyses were performed using GraphPad Prism version 7 (GraphPad Software, San Diego, CA, USA).

#### Western blot

Differences between groups were tested for significance using unpaired t-tests.

#### qPCR

Differences between groups were tested for significance using two-way ANOVA. Holm– Sidak’s multiple comparisons *post hoc* tests were performed, when appropriate.

#### Correlation between RNA-seq and ChIP-seq data-sets

Spearman correlation was run on the datasets. The results were considered significant when p was <.05.

#### Locomotor activity

differences between total activity and circadian rhythmicity were tested using a two-way RM ANOVA.

All data are represented as the mean ± SEM unless otherwise stated. N refers to single animals unless otherwise stated. In the figures *p<.05; **p<.001, ***p<.0001. Error bars represent SEM.

### Data availability

RNA-seq and ChIP-seq data are available on the GEO database: accession number GSE168725

## Supporting information

Supplementary Informations

## ACKNOWLEDGMENTS

We thank Manuel Tongiani, Francesca Damiani and Matteo Caldarelli for their help with the experiments. Special thanks to Vania Liverani and Antonella Calvello (Scuola Normale Superiore) for technical assistance in the lab. We thank Prof. Concetta Morrone and Prof. Paola Binda (University of Pisa) for their insightful comments.

This research was supported by H2020-MSCA-IF-2016 749697 GaMePLAY, University of Pisa PRA-2020 to PT. The work of SC, MS, CM, and PB was in part supported by NIH grant GM123558 to PB.

## Author Contributions

SCo and LL performed all the experiments and analysed the data. SCo prepared the figures. SCh performed the ChIP-seq and RNA-seq analysis. FF and SR performed the Beta-hydroxybutyrilome analysis. MS performed the RNA seq analysis. FR helped with GO analysis. RM helped with correlations between ChIP-seq and RNA-seq. CM helped with ChIP-seq data alignment and analysis. PB supervised the RNA-seq and ChiP-seq data analysis. PT conceived the project and performed the experiments. TP and PT supervised the project and wrote the manuscript.

