## Supplementary Informations for "Brain histone beta-hydroxy-butyrylation couples metabolism with gene expression"

### SUPPLEMENTARY MATERIALS

#### RNA extraction

Occipital cortex samples were homogenized in phenol/guanidine-based QIAzol Lysis Reagent (Qiagen #79306). Chloroform was added and the samples were shacked for 15 s. The samples were left at 20–24°C for 3 min and then centrifuged (12000*xg*, 20 min, 4 °C). The upper phase aqueous solution, containing RNA, was collected in a fresh tube and the RNA was precipitated by the addition of isopropanol. Samples were mixed by vortexing, left at RT for 10 min and then centrifuged (12000*xg*, 15 min, 4 °C). The supernatant was discarded and the RNA pellet was washed in 75% ethanol by centrifugation (7500*xg*, 5 min, 4 °C). The supernatant was discarded and the pellet was left to dry for at least 15 min; then, it was resuspended in RNAse free water.
Due to the small amounts of tissue, samples containing the suprachiasmatic nucleus of the hypothalamus were homogenized in phenol/guanidine-based QIAzol Lysis Reagent (Qiagen #79306) and then processed using the miRNeasy Mini Kit (Qiagen # 217004).
RNA concentration was determined by Nanodrop Spectrophotometer (Thermoscientific 2000 C). RNA quality was analyzed through a gel running (1% agarose). Total RNA was reverse transcribed using QuantiTech Reverse Transcription Kit (Qiagen # 205311). Gene expression was analyzed by real-time PCR (Step one, Applied Biosystems), using PowerUp SYBR Green Master Mix (Thermo Fisher #A25742). The primers for gene expression were designed by Primer 3 software (v. 0.4.0) and reported in the following table:

#### Primer Table

| | |
|---|---|
| mBmal1 FOR | 5' GCAGTGCCACTGACTACCAAGA 3' |
| mBmal1 REV | 5' TCCTGGACATTGCATTGCAT 3' |
| mCry2 FOR | 5' CAACACAGGCCCCAGAGCACTATC 3' |
| mCry2 REV | 5' TCAGGAGTCCTTGCTTGCTGGCTC 3' |
| mCry1 FOR | 5’ CAGACTCACTCACTCAAGCAAGG 3’ |
| mCry1 REV | 5’ TCAGTTACTGCTCTGCCGCTGGAC 3’ |
| mReverb-a FOR | 5' AGGCTGCTCAGTTGGTTGTT 3' |

| | |
|---|---|
| mReverb-a REV | 5' CTCCATCGTTCGCATCAATC 3' |
| mPer1 FOR | 5’ ACCAGCGTGTCATGATGACATA 3’ |
| mPer1 REV | 5’ GTGCACAGCACCCAGTTCCC 3’ |
| mRor-a FOR | 5' ACCGTGTCCATGGCAGAAC 3' |
| mRor-a REV | 5' TTTCCAGGTGGGATTTGGAT 3' |
| mClock FOR | 5' ACCACAGCAACAGCAACAAC 3' |
| mClock REV | 5' GGCTGCTGAACTGAAGGAAG 3' |

Quantitative values for cDNA amplification were calculated from the threshold cycle number (Ct) obtained during the exponential growth of the PCR products. The threshold was set automatically by the Step one software. Data were analyzed by the ΔΔCt methods using 18S rRNA to normalize the cDNA levels of the transcripts under investigation.

**Protein extraction and Western blot**

For total protein extracts, liver and cerebral cortex samples were homogenized in modified RIPA buffer (50 mM Tris pH8, 150mM NaCl, 5mm EDTA, 15mM MgCl2, 1% NP40) plus protease inhibitors. The samples were sonicated for one minute on ice (10 secs on/10 secs off) and centrifuged for 15 min at 14000g 4°C. The supernatant was recovered and the protein concentration was determined by Bradford assay (Biorad #5000006) using a Nanodrop Spectrophotometer (Thermoscientific 2000 C).

8% SDS-PAGE was performed to check bhb-Lysine, phospho(Ser235-236) S6, total S6, phospho (Ser473)AKT and total AKT levels. 15% SDS-PAGE was performed to analyse H3 and total H3K9-bhb. The samples were blotted onto nitrocellulose membranes (Biorad) and blocked in 5% BSA in Tris-buffered saline (TBS) for 1 hour at room temperature RT. The nitrocellulose membrane was incubated at 4°C overnight with the following antibodies: anti-phospho(Ser235-236) S6 (Cell signalling Technology #2211S) 1:1000, anti-S6 (Cell signalling Technology #2217) 1:1000, and anti-phospho (Ser473) AKT 1:1000 (Cell signalling technology #9271), AKT 1:1000 (Cell signalling technology #9272), bhb-Lysine 1:2000 (PTM Biolabs #1201), H3K9-bhb 1:2000 (PTM Biolabs #1250), H3 1:5000 (Abcam #1791), beta-tubulin 1:3000 (Sigma #T4026). Blots were then washed 3 times in TTBS for thirty minutes, incubated in HRP conjugated anti-mouse or anti-rabbit diluted (1:8000) in 2.5% BSA in TTBS for one hour at RT. The membranes were then rinsed three times in TTBS and incubated in enhanced chemiluminescent substrate (Millipore) and acquired through a Chemidoc XRS instrument. Bands densitometry was analyzed through ImageJ software.

**Beta-hydroxybutyrilome analysis**

Cerebral cortex samples were homogenized in 50 mM Tris-HCl, 120 mM NaCl, 1% sodium deoxycholate, briefly sonicated on ice, and centrifuge for 15 min at 14000g 4°C. The supernatant was recovered and the protein concentration was determined by Bradford assay (Biorad) using a Nanodrop Spectrophotometer (Thermoscientific 2000 C). For each condition, 50 µg of proteins were reduced with dithiothreitol (10 mM, for 30 min, at 65°C) and alkylated using iodoacetamide (22 mM, 30 min, room temperature) in dark conditions. Protein digestion was performed using trypsin (w/w ratio 1:50) at 37 °C for 16 hours. Samples were incubated with 1% trifluoroacetic acid (Sigma-Aldrich) for 45 min at 37°C to quench trypsin reaction and to remove sodium deoxycholate by acid precipitation. Samples were centrifuged at 16,000 x g for 10 min and subsequently desalted with Mobicol spin columns equipped with 10 µm pore size filters and filled with VersaFlash C18 spherical 70 Å silica particles (Sigma-Aldrich). Peptides were lyophilized and consequently dissolved in 2% acetonitrile / 0.1 formic acid to achieve a final peptide concentration of 1µg/µL before liquid chromatography tandem mass spectrometry (LC-MS/MS) analysis. The equivalent of 5 µg per sample were directly loaded with an Eksigent expert™ microLC 200 system (Eksigent, AB Sciex, Framingham, MA, USA) and acquired using a 5600+ TripleTOF mass spectrometer (AB Sciex). After loading, peptides were separated on a Jupiter 150 x 0.3 mm, 4 µm 90 Å capillary column with a gradient from 2 to 35% buffer B (acetonitrile / 0.1 formic acid) in 40 min at a flow rate of 5 µL/min. LC column was directly interfaced with a DuoSpray™ ESI ion source operating at 5.5 kV for peptide ionization. Samples were analyzed with an information-dependent acquisition (IDA) tandem mass spectrometry method based on a MS1 survey scan from which the 20 most abundant precursor ions were selected for subsequent CID fragmentation to generate MS2 spectra. MS1 survey and MS2 scans were acquired with a resolving power of 30000 and 25000 and over a mass range of 250 – 1250 m/z and 150 -1500 m/z, respectively. Isolation width for precursor ion selection was set at 0.7 m/z on a Q1. The accumulation time was set to 250 milliseconds for MS1 scans while 100 milliseconds for MS2 scans. Charge states of 1+ were excluded from ion selection. Rolling collision energy with a collision energy spread across 5 eV and background subtraction were enabled to achieve the optimal fragmentation according to m/z ratio and charge state and to increase sensitivity. Peak areas of peptides associated to unique proteins were summed together to achieve total protein abundance values that were normalized through the total ion current (TIC) extracted from the full MS1 survey scan acquisition for each run. Proteins were considered to be significantly different with a p-value lower than 0.05 and a fold change (|FC|) > 1.5.

**Chromatin Immunoprecipitation (ChIP)**

Minced frozen cortices were crosslinked with 1% formaldehyde for 10 min followed by Glycine (0.125M final concentration) at room temperature for 10 min. After homogenizing tissue pellets in PBS, 1 ml of lysis buffer was added. Samples were sonicated by Bioruptor (15 cycles, every cycle: 30 sec ON / 30 sec OFF, power high) to generate 200-500 base pairs fragments and centrifuged at 14000g at 4°C. Supernatants were diluted in dilution buffer (1.1% Triton X100, 1.2 mM EDTA, 16.7 mM Tris-HCl, 167 mM NaCl), and incubated with the following primary antibody 3ug H3K9bhb (PTM Biolab #1250) overnight at 4°C. To monitor the specificity of ChIP assays, samples were also immunoprecipitated with a specific-antibody isotype matched control immunoglobulin (IgG). Protein-G beads (Invitrogen #10004D) were added to the supernatant and incubated for 2 hrs at 4°C and centrifuged. Beads were recovered, washed in low salt buffer, high salt buffer, LiCl buffer, followed by washing in TE for three times. Elution buffer (300 mM NaCl, 0.5% SDS, 10 mM Tris-HCl, 5mM EDTA) was added to the washed beads, treated with RNase at 37°C for 2 hrs and Proteinase K at 65°C overnight. Equal amount of Phenol-Chloroform-Isoamyl Alcohol was added to the samples and the aqueous phase was recovered. DNA was precipitated by adding 100% Ethanol, NaAc and glycogen and kept at -20°C overnight. Samples were centrifuged at 14000g for 30 min at 4°C and washed with 70% ethanol followed by centrifugation at 14000g for 15 min at 4°C.

**Gene ontology (GO) terms analysis and Functional network generation**

Gene ontology analysis was performed using Database for Annotation, Visualization and Integrated Discovery (DAVID) v6.7, using genomic background; and the Kyoto Encyclopedia of information-dependentGenes and Genomes (KEGG) pathway and GO Biological Processes were chosen for gene clustering. To confirm the results, the CAMERA approach (1, 2), available through EnrichmentBrowser (3), was also used for gene-set enrichment analysis after gene-ranking based on differential expression log fold changes. Cytoscape 3.7.2 software was used to visualize complex networks of interactions in the data-sets, and integrate them with functional annotations.

### SUPPLEMENTARY FIGURES LEGEND

**Fig. S1**

(**a**) Western blot analysis of lysine beta-hydroxybutyrylation in the liver of F and AL mice using a specific panK9-bhb antibody. (**b**) Western blot analysis of histone K-bhb in the liver of F or AL using a specific H3K9-bhb antibody (top panel), H3 loading control (middle panel), histograms representing the normalized ratio of H3K9-bhb /H3 in F condition compared to AL (low panel) (AL N=4, F N=4, unpaired 2-tailed t-test, $t_6$=3.203 p=0.0185). Error bars represent SEM.

**Fig. S2**

(a) Pie chart of genome-wide distribution of H3K9-bhb enriched loci in F with respect to AL (p<0.05) in the liver, data from (4) reanalyzed.

**Fig. S3**

Interaction network of DE genes using Cytoscape.

**Fig. S4.**

Gene enrichment analysis of the KEGG pathway mmu04710 “Circadian rhythms” on the whole RNA-seq data set using CAMERA. (**a**) Only transcripts DE: BH<0.01. (**b**) All transcripts.

**Fig. S5**

(**a**) Western blot analysis of phospho (p) (Ser473) AKT in the occipital cortex of F or AL (top panel), AKT (middle panel), histograms representing the normalized ratio of p(Ser473) AKT/AKT in F condition compared to AL (low panel); (AL N=5, F N=6, unpaired 2-tailed t-test, $t_9$=3.893 p=0.0037). (**b**) Western blot analysis of phospho (p) (Ser235-236) S6 in the occipital cortex of F or AL (top panel), S6 (middle panel), histograms representing the normalized ratio of pS6/S6 in F condition compared to AL (low panel); (AL N=5, F N=6, unpaired 2-tailed t-test, $t_9$=5.55 p=0.0004). Error bars represent SEM.

**Fig. S6**

(**a**) Schematic coronal section of the SCN showing the site of the gene expression analysis. (**b**) Experimental timeline. (**c**) Quantitative real-time PCR analysis of genes belonging to the inhibitory limb of the core-clock machinery (N=4 per condition, per time-point). Per1: two-way ANOVA time*treatment, interaction $F_{3,24}$=10.55 p=0.0001, post-hoc Sidak, AL vs F (ZT22) $t_{24}$=6.526 p<0.0001; Cry1: two-way ANOVA time*treatment, interaction $F_{3,24}$=5.241 p=0.0063, post-hoc Sidak, AL vs F (ZT4) $t_{24}$=2.808 p=0.0384, AL vs F (ZT22) $t_{24}$=3.907 p=0.0027; Cry2: two-way

ANOVA time*treatment, interaction $F_{3,24}$=13.23 p<0.001, post-hoc Sidak, AL vs F (ZT22) $t_{24}$=8.377 p<0.001, (ZT22) $t_{24}$=2.769 p=0.042; Reverb-a, two-way ANOVA time*treatment, interaction $F_{3,24}$=9.149 p=0.003, post-hoc Sidak, AL vs F (ZT22) $t_{24}$=5.371 p<0.001. **d**) Quantitative real-time PCR analysis of genes belonging to the transcriptional activation limb of the core-clock machinery (N=4 per condition, per time-point). Bmal1: F, two-way ANOVA time*treatment, interaction $F_{3,24}$=2.546 p=0.0798; Clock: two-way RM ANOVA time*treatment, interaction $F_{3,24}$=1.618 p=0.2127; RORa: F, two-way RM ANOVA time*treatment, interaction $F_{3,24}$=1.105 p=0.3673). Error bars represent SEM.

**Fig. S7**

**a**) Experimental timeline. **b**) Relative power spectra of the locomotor activity in AL and FAST mice (N=6 per group; bold lines represent the mean, shaded areas represent the SEM) in three epochs: before fasting (Pre-fast), first five days of refeeding (T1) and second 5 days of refeeding (T2). **c**) Total locomotor activity measured as the number of beam breaks of AL and FAST mice during the light phase (top) and the dark phase (bottom) for each day. The area shaded in gray represents the fasting period. (Light phase activity: two-way RM ANOVA time*treatment; time: F1,10=0.287 p=0.604, treatment: F1,10=0.082 p=0.781, interaction: F1,10=0.012 p=0.915. Dark phase activity: two-way RM ANOVA time*treatment; time: F1,10=16.72 p=0.002, treatment: F1,5=0.704 p=0.421, interaction: F1,5=1.247 p=0.290) ***d)*** Weight change (%) before and after 48h fasting, and during the refeeding phase at two different timepoints : 5 days after the refeeding (T1), 10 days after the refeeding (T2) (AL N=4, F N=4, Mixed-effects analysis time*treatment, interaction F3,10=22.3 p<0.0001, post-hoc Sidak, AL vs F (post-fasting) t16=8.956 p<0.0001;). Error bars represent SEM.
